# Brain-heart interactions are optimized across the respiratory cycle via interoceptive attention

**DOI:** 10.1101/2022.04.02.486808

**Authors:** Andrea Zaccaro, Mauro Gianni Perrucci, Eleonora Parrotta, Marcello Costantini, Francesca Ferri

## Abstract

Respiration and heartbeat continuously interact within the living organism at many different levels, representing two of the main oscillatory rhythms of the body and providing major sources of interoceptive information to the brain. Despite the modulatory effect of respiration on exteroception and cognition has been recently established in humans, its role in shaping interoceptive perception has been scarcely investigated so far.

In two independent studies, we investigated the effect of spontaneous breathing on cardiac interoception by assessing the Heartbeat Evoked Potential (HEP) in healthy humans. In Study 1, we compared HEP activity for heartbeats occurred during inhalation and exhalation in 40 volunteers at rest. We found higher HEP amplitude during exhalation, compared to inhalation, over fronto-centro-parietal areas. This suggests increased brain-heart interactions and improved cortical processing of the heartbeats during exhalation. In Study 2, we tested the respiratory phase-dependent modulation of HEP activity in 20 volunteers during Exteroceptive and Interoceptive conditions of the Heartbeat Detection (HBD) task. In these conditions, participants were requested to tap at each heartbeat, either listened to or felt, respectively. Results showed higher HEP activity and higher detection accuracy at exhalation than inhalation in the Interoceptive condition only. These effects were positively correlated, suggesting a link between optimization of both cortical processing of cardiac signals and perception of heartbeats across the respiratory cycle. Direct comparisons of Interoceptive and Exteroceptive conditions confirmed stronger respiratory phase-dependent modulation of HEP and accuracy when attention was directed towards the interoceptive stimuli.

Overall, we provide data showing that respiration shapes cardiac interoception at the neurophysiological and behavioural levels. Specifically, exhalation may allow attentional shift towards the internal bodily states.

## Introduction

Respiration and heartbeat are inextricably interconnected. Together, they produce two of the dominant oscillatory rhythms of the organism and represent major sources of interoceptive information (Chen et al., 2021; Khalsa et al., 2009; Weng et al., 2021). Interoception has been commonly defined as the process by which the brain receives, elaborates and interprets signals originating from the peripheral organs, continuously updating the conscious and (mostly) unconscious representations of the physiological condition of the body (Berntson and Khalsa, 2021; Craig, 2003; Critchley et al., 2004). However, within the field of interoception research, respiratory and cardiac signals processing have been studied mainly separately so far. In addition, there is no question that cardiac interoception has attracted most of the attention in the field. Since the Schandry’s first proposal (Schandry, 1981), various behavioural tasks have been suggested to assess the individual interoceptive accuracy, as the ability to voluntarily focus on one’s own heart and correctly report its beating. Importantly, research has shown that higher sensitivity to cardiac signals supports the capacity to regulate emotions and behaviour (Dunn et al., 2007; Herbert et al., 2011, 2012; Herbert and Pollatos, 2014; Wiens, 2005), while it is negatively associated to the susceptibility to mental health problems (de la Fuente et al., 2019; Lutz et al., 2019; Schulz et al., 2015; Yoris et al., 2017). However, heartbeat sensations remain for the most part outside of the field of awareness, and methodological limitations of cardiac interoceptive tasks have emerged (Brener and Ring, 2016; Garfinkel et al., 2022).

More recently, researchers have increasingly focused on a more objective, electrophysiological index of the cortical processing of single heartbeats, the so-called Heartbeat Evoked Potential (HEP). The HEP is an Electroencephalographic (EEG) event-related potential, time-locked to participants’ Electrocardiogram (ECG) R-peak or T-peak (Pollatos and Schandry, 2004). Physiological pathways underlying the HEP are currently under study, but they mainly involve signals originating from baroreceptor activity. Baroreceptors are stretch receptors located near the aortic arch and the carotid arteries, whose discharge activity is time-locked to cardiac systole and is driven by rhythmic changes in arterial blood pressure (Park and Blanke, 2019a). Baroreceptor-mediated information is sent upstream via the vagus nerve to the nucleus of the solitary tract in the brainstem and to the ventromedial posterior nucleus of the thalamus. Cardiac interoceptive information is finally elaborated in the brain at the level of the insula, amygdala, anterior cingulate, and somatosensory cortices, which represent the main cerebral sources of the HEP activity (Babo-Rebelo et al., 2016; Canales-Johnson et al., 2015; Kern et al., 2013; Park and Tallon-Baudry, 2014). Importantly, HEP activity increases when individuals voluntarily orient their attention to the heartbeat (García-Cordero et al., 2017; Mai et al., 2018; Petzschner et al., 2019; Salamone et al., 2018; Villena-González et al., 2017), and is also positively associated to their accuracy in detecting heartbeats during cardiac interoceptive tasks (Canales-Johnson et al., 2015; Marshall et al., 2017; Pollatos et al., 2005).

In line with the inextricable interconnection between heartbeat and respiration, separate studies have independently shown that both HEP and cardiac interoceptive accuracy are modulated by respiration. For instance, one study from Baumert et al. (2015) assessed respiratory phase-dependent (inhale vs. exhale) HEP peak amplitude in children with sleep disordered breathing during REM sleep. The authors found decreased peak HEP amplitude during exhalation in the sleep disordered breathing group. Moreover, MacKinnon and colleagues (2013) observed increased peak HEP amplitude over central EEG electrodes during “resonant breathing” (i.e., breathing at a rate of 6 breaths per minute), compared to spontaneous breathing. Finally, two recent studies (Smith et al., 2020, 2021) consistently showed that breath-holding can improve participants’ cardiac interoceptive accuracy, as assessed with a modified version of the Heartbeat Detection (HBD) task. However, all these studies investigated the effect of respiration on HEP changes and interoceptive accuracy separately and in a “perturbated” physiological setting, that is during systemic alteration of respiratory activity (REM sleep, slow breathing, and breath-hold). A systematic and multilevel investigation of the complex neuro-cardio-respiratory interactions (Corcoran et al., 2018) in an “ecological” setting is still lacking. This would allow the simultaneous characterization of the effects of spontaneous breathing on both HEP and interoceptive accuracy, as well as their possible relationship.

Spontaneous breathing occurs mostly outside of the field of awareness and comprises a phase of active inspiration, that involves the contraction of the diaphragm and the external intercostals muscles, and a phase of passive expiration due to their subsequent relaxation. However, unlike the heartbeat, breathing can be easily accessed consciously and voluntarily controlled in its depth and frequency (Del Negro et al., 2018; Feldman et al., 2013). Respiration represents one of the most salient conscious forms of interoception. Despite this, it has received relatively low scientific interest until very recently, when increasing evidence has been accumulating about respiratory phase-dependent changes of perception and cognitive-emotional functions. For example, Zelano et al. (2016) found increased emotional recognition and episodic memory encoding and retrieval during inhalation, as compared to exhalation. Furthermore, phase-locking of stimulus onset to inhalation increased participant’s near-threshold somatosensory perception (Grund et al., 2021) visuospatial recognition abilities (Kluger et al., 2021; Perl et al., 2019), and memory performance (Huijbers et al., 2014). Differently, self-initiated motor actions (Park et al., 2020), and trace eyeblink conditioning learning (Waselius et al., 2019) were more frequent during exhalation, while the exhalation-to-inhalation phase transition improved recognition memory performance (Nakamura et al., 2018). Based on this and other evidence, within the framework of predictive coding theories of brain functioning (Clark, 2013; Friston, 2010), respiration has been recently interpreted as a form of “active sensing” (Allen et al., 2021; Boyadzhieva and Kayhan, 2021; Corcoran et al., 2018). Active sensing indicates that the rhythmic neuronal excitability changes induced by inhalation adaptively align bottom-up sensory information with top-down predictive streams (Corcoran et al., 2018), amplifying incoming sensory information from the environment, namely, exteroceptive information (Tort et al., 2018). Hence, the question arises of whether spontaneous respiration plays a role in shaping the processing of interoceptive information generated from inside the body as well.

In particular, the role of the respiratory phases assumes a special psychophysiological interest in the context of cardiac interoception for the following reasons: first, heartbeat and respiration are deeply linked at the body level, given the coupling between respiratory cycle and baroreceptor activity. This interaction is commonly observed during respiratory phase-dependent heart rate changes known as Respiratory Sinus Arrhythmia (RSA, Brecher and Hubay, 1955). Second, heartbeat and respiration share similar interoceptive pathways, both reaching the central nervous system through the vagus nerve at the level of the anterior and posterior insula, hippocampus, precuneus, somatosensory, and cingulate cortices (Farb et al., 2013; Wang et al., 2019), suggesting the possibility that respiratory and cardiac interoceptive signals interact also at the brain level, with possible effects on the individual interoceptive accuracy.

Therefore, the aim of the present work is twofold: i) to investigate and characterize respiratory phase-dependent modulations of neural responses to single heartbeats, as measured by HEP activity at rest; and ii) to relate respiratory phase-dependent HEP modulations to cardiac interoceptive attention. To fulfil these objectives, we performed two independent studies, simultaneously recording participant’s EEG, ECG, and respiratory activity. In Study 1, we tested 40 healthy volunteers during a resting-state condition, while in Study 2 we tested 20 healthy volunteers during the performance of the HBD task, which requires to focus attention on the heart and tap a button in synchrony with each heartbeat.

## Materials and methods

### Ethics Statement

The studies were approved by the Institutional Review Board of Psychology, Department of Psychological, Health and Territorial Sciences, “G. d’Annunzio” University of Chieti-Pescara (Protocol Number 44_26_07_2021_21016), in compliance with the Italian Association of Psychology and the Declaration of Helsinki guidelines and its later amendments. All subjects signed a written informed consent.

### Study 1 – Resting-state condition

#### Participants

Forty healthy volunteers with normal or corrected-to-normal vision (29 females; two left-handed; age: 26.67 ± 4.56 years [mean ± SD]) took part in the study. We estimated the sample size through the G*Power 3 software (v3.1.9.7; Faul et al., 2007) based on the results of a recent meta-analysis on HEP activity (Coll et al., 2021). We estimated a medium effect size of Cohen’s d = 0.5, set the significance level to alpha = 0.05, and the desired power at 0.80 (estimated sample size = 34). The inclusion of each volunteer was based on the following criteria: i) no personal or family history of neurological, psychiatric, or somatic disorders; and ii) not having taken any drug acting on the central nervous system in the previous week. One participant was excluded from EEG analysis because of excessive movement-related artifacts.

### Experimental procedure

Participants were asked to rest for 10 minutes with eyes open while watching at a fixation cross at the centre of a computer screen (Raichle et al., 2001). No specific instruction on breathing was given, and participants were blind to the experimental goals. EEG, ECG, and respiratory signals were simultaneously recorded throughout the session. Before starting the session, participants had to verbally confirm that they could not feel their heartbeat through the respiratory belt.

### Electrophysiological recordings

EEG signal was recorded from 64 scalp electrodes using a BrainAmp EEG acquisition system (BrainCap MR, BrainVision, LLC), according to the international 10-20 system. The midfrontal electrode (FCz) was used as the reference and the inion electrode (Iz) as the ground. Electrode impedance was kept below 10 kΩ for all channels. ECG data were obtained from three ECG electrodes, two placed over the left and right clavicles, and the ground located on the right costal margin (BIOPAC Systems, Inc). Another ECG electrode, that was integrated in the EEG net, was placed on the left breast, serving as a backup. Respiratory activity was recorded via a respiratory belt positioned around the chest (respiratory transducer TSD201, BIOPAC Systems, Inc). All signals were recorded with a sampling rate of 2 kHz; band-pass filtering from 0.016 to 250 Hz was applied, along with 50 Hz notch filtering.

### Electrophysiological data pre-processing

ECG and respiratory signals were high-pass filtered (0.1 Hz) to remove baseline fluctuations. A low-pass filter at half the resampling frequency (i.e., 128 Hz) was applied to the data to avoid aliasing effects, then signals were down-sampled to 256 Hz. R-peaks were extracted from the ECG using the Pan-Tompkins algorithm (Pan and Tompkins, 1985; Sedghamiz, 2014), and mis-detected peaks were corrected using a point process model (Citi et al., 2012). The obtained RR-interval data set (tachogram) was processed using Kubios Oy free software (v3.4.3, Tarvainen et al., 2014) and a set of Heart Rate Variability (HRV) features of interest were extracted. Time-domain parameters, such as Heart Rate (HR), and frequency-domain parameters, such as power in High-Frequency band (HF −0.15-0.4 Hz) log value (Malik et al., 1996), Heart Rate Variability (HRV) total power, and Low-Frequency/High-Frequency ratio (LF/HF) (Supplementary Material 1) were computed. Respiratory signals were processed using BreathMetrics (Noto et al., 2018) toolbox algorithms, to extract inhale and exhale onsets and offsets, as well as the set of respiratory features of interest, such as breathing rate, average inhale duration, average exhale duration, Inhalation/Exhalation (I/E) ratio (Supplementary Material 1). Inhale and exhale onsets and offsets were then visually inspected, and noisy breathing cycles were manually rejected. A respiratory cycle was defined as starting from the beginning of an inhalation and ending at the beginning of the next inhalation.

EEG data were pre-processed offline using EEGLAB (v2021.1; Delorme and Makeig, 2004) toolbox algorithms running on a MATLAB environment (R2021a, MathWorks Inc.). After all individual blocks were concatenated, signals were down-sampled to 256 Hz. Before resampling, a low-pass filter at half the resampling frequency (i.e., 128 Hz) was applied to the data to avoid aliasing effects. Datasets were then filtered using a Hamming windowed FIR filter (0.5-40 Hz). Signals were visually inspected for the removal of artefacts and the detection of noisy channels. Bad segments were manually rejected. Noisy EEG channels were then removed and interpolated using their neighbouring channels (Al et al., 2021; Junghöfer et al., 2000). Rejected channels were generally few (~5%, depending on the EEG recording). Retained signal was submitted to Independent Component Analysis to visualize and manually remove sources of heartbeat, ocular, and muscle artifacts (FastICA algorithm; Hyvärinen, 1999). Particular attention was given to cardiac field artifact (CFA), by visually selecting the components whose activities followed the time course of R-peak and/or T-peak of the ECG (Al et al., 2020, 2021). EEG signals were finally re-referenced to the average of all channels.

### HEP analysis

HEP was analysed using ERPLAB toolbox algorithms (v8.30; Lopez-Calderon and Luck, 2014). HEP was computed on EEG signals time-locked to the T-peak of the ECG (Babo-Rebelo et al., 2019; Babo-Rebelo et al., 2016; Park et al., 2014). ECG T-peak positions were identified using the HEPLAB toolbox (Perakakis, 2019). Automatic detection of T-peaks was then followed by visual inspection and manual correction (Al et al., 2021). EEG data were epoched and baseline-corrected between −100 and 0 ms, using the T-peak event as temporal reference (epoch length: −100 to 350 ms after T-peak) (Park et al., 2014). The time window of interest for the statistical analysis was set to 80-350 ms, coincident with cardiac relaxation, when the CFA is minimum (Babo-Rebelo et al., 2019; Babo-Rebelo et al., 2016; Dirlich et al., 1997). Additionally, we rejected all epochs in which the signal recorded at any channel exceeded a threshold of 100 μV (Blankenship et al., 2018; Villena-González et al., 2016), using a moving window peak-to-peak threshold function implemented in ERPLAB (moving window size: 200 ms; step size: 100 ms) (Lopez-Calderon and Luck, 2014). The number of rejected epochs was less than 1%. Artifact-free epochs were assigned to “inhale” if the respective T-peak fell within the inhale onset/inhale offset time range, and to “exhale” if the T-peak fell within the exhale onset/exhale offset time range. HEP corresponding to inhale and exhale were computed for each participant by averaging EEG epochs assigned to the respective respiratory phase. We analysed HEP epochs occurring exclusively during inhalation or exhalation, that is, excluding those occurring across both respiratory phases, to avoid uncontrolled effects due to inhalation or exhalation onset (e.g., respiratory-related evoked potentials; Davenport et al., 2007; Webster and Colrain, 2000). Finally, we excluded epochs including T-peaks followed by an R-peak by less than 350 ms, to avoid overlap between the HEP activity and the following R-peak residual CFA (Babo-Rebelo et al., 2019).

HEP differences between respiratory phases were statistically assessed in the EEG artifact-free time window 80–350 ms after the T-peak, using a cluster-based permutation t-test implemented in the FieldTrip toolbox (Oostenveld et al., 2011). First, paired t-values were calculated by comparing the “inhale” and “exhale” phases in the 80-350 ms time window. Clusters were then formed by pooling together all adjacent spatiotemporal data points with a p-value below 0.05 within the time window of interest. The t-statistic of each cluster was obtained by summing all t-values within the cluster. Then, a randomized null distribution of cluster-level t-statistics was obtained with a Monte Carlo permutation procedure that randomly shuffled the “inhale” and “exhale” phase labels 10000 times, and entered into the null distribution the largest obtained cluster-level statistic for each randomization. Finally, statistical significance was calculated by comparing the experimentally observed cluster-level statistics with the randomly-generated null distribution. This procedure inherently corrects for multiple comparisons in time and space, and clusters with a corrected p-value below 0.05 were considered significant (two-tails). We did not use any a priori spatial or temporal region of interest for the comparison between inhale and exhale HEP, hence including the entire EEG sensor space and epoch time window (80-350 ms) (Petzschner et al., 2019).

### Source analysis

A standard structural T1-weighted MRI template (ICBM152) (Fonov et al., 2009) was used to estimate the neural sources of the EEG signals within the BrainStorm toolbox (v3.210416; Tadel et al., 2011). Lead field matrix were computed with a 3-shell Boundary Element Model using the OpenMEEG (Gramfort et al., 2010; Kybic et al., 2005) toolbox. Then, neural sources were estimated following the Minimum Norm Estimation method using sLORETA normalization (Pascual-Marqui, 2002), by keeping constrained current dipole orientations (i.e., normally oriented) with respect to the cortical surface. For the statistical test of the neural sources of differential HEP amplitudes among respiratory phases (inhale vs. exhale), we used cluster-based statistics in the source space (1000 randomizations) from the FieldTrip toolbox (Oostenveld et al., 2011).

### Additional cardiorespiratory analyses during resting-state

To investigate the possible explanatory role of cardiac and respiratory physiology on HEP activity among respiratory phases, we performed the following additional analyses. First, to exclude confounding influence of the CFA, we tested for differences in the ECG signal amplitude between the inhale and exhale phases (Petzschner et al., 2019). We submitted the averaged ECG signal time-locked to the T-peak (epoch length: −100 to 350 ms) from the inhale and exhale phases to a repeated-measures, two-tailed t-test for all time points within the time window of observed significant HEP differences (Groppe et al., 2011), corrected with the False Discovery Rate (FDR, Benjamini and Yekutieli 2001) procedure. In addition, we calculated ΔHEP for each participant by subtracting the mean HEP value during inhale from the mean HEP value during exhale in the significant time window, averaged within the cluster of significant electrodes. Next, we computed ΔECG by subtracting the mean ECG value during inhale from the mean ECG value during exhale in the significant time window. Finally, we tested for any relationship between cardiorespiratory activity and HEP changes across participants using Pearson’s correlation analyses relating mean HEP changes among respiratory phases (ΔHEP) to both changes in the ECG (ΔECG) and participants’ cardiac and respiratory features of interest (Supplementary Material 1).

### Study 2 - Heartbeat Detection Task

#### Participants

Twenty healthy volunteers with normal or corrected-to-normal vision took part in the study (14 females; two left-handed; mean age: 25.21 ± 2.64 years [mean ± SD]). Sample size was estimated through G*Power 3 (v3.1.9.7; Faul et al., 2007) based on the results of a previous study investigating HBD task accuracy (Fittipaldi et al., 2020). We estimated a medium/high effect size of Cohen’s d = 0.8, set the significance level to 0.05, and the power at 0.80 (estimated sample size = 15). Inclusion criteria were the same as in Study 1. One participant was excluded from EEG analysis because of excessive movement-related artifact.

### Experimental procedure

Following a brief training, participants were asked to perform the two conditions of a validated HBD task (Fittipaldi et al., 2020; García-Cordero et al., 2017; Yoris et al., 2017). In the first condition, named Exteroceptive Condition (EC), participants were presented with digitally constructed heartbeat sounds and instructed to press a button in synchrony with it using their dominant hand. They were given the following instructions: “You will hear the beating of a heart. Tap the button with your dominant hand as soon as you hear each heartbeat. Please avoid anticipated responses by guessing the recorded heart rhythm”. The EC consisted of 4 blocks, each lasting 2.5 minutes, for a total of 10 minutes. In two blocks the heartbeats were presented at a regular frequency (60 bpm), while in the rest of the blocks the heartbeats were presented with irregular heartbeat intervals. In the second condition, named Interoceptive Condition (IC), participants were asked to focus their attention on their heart and tap a button in synchrony with their own heartbeats, in the absence of any external cues. Instructions were as follows: “Now, you must follow the beating of your own heart by tapping a button with your dominant hand for every beat you feel. You should not guide your responses by checking your arterial pulse in your wrists or neck. If you are unable to feel these sensations, you should appeal to your intuition trying to respond whenever you think your heart is beating” (Fittipaldi et al., 2020). The IC consisted of 4 blocks lasting 2.5 min each, for a total of 10 minutes. As in Study 1, during both conditions, participants kept their eyes open and watched a fixation cross at the centre of the monitor. No specific instructions on breathing were given. EEG, ECG, and respiratory signals were simultaneously recorded throughout IC and EC.

### Electrophysiological recordings, signals pre-processing, and HEP analysis

We applied the same analysis pipeline of Study 1 for recording EEG and cardiorespiratory signals, as well as for EEG data pre-processing and HEP analysis.

### Task accuracy assessment and comparison

Participants’ interoceptive accuracy was assessed by calculating the interoceptive accuracy index, based on previously reported standard procedures (Fittipaldi et al., 2020; García-Cordero et al., 2017; Yoris et al., 2017). Correct answers were estimated by time-locking participant’s tapped responses with a corresponding time window around each R-peaks registered by the ECG. To control for HR differences between participants in the IC, we considered three different time windows depending on the participant’s HR. Unlike canonical procedure, which calculates the time windows of accurate response depending on mean HR, we relied on the participants’ instantaneous HR. Instantaneous HR (bpm) was calculated for each R-peak based on the time from the current R-peak of the heart wave to the R-peak point of the next wave. This allowed us to control for ongoing HR changes during the task, in particular, for those coupled with the respiratory cycle (i.e., RSA). We thus considered 750 ms after the R-peak for an instantaneous HR less than 69.76 bpm; 600 ms after the R-peak for an instantaneous HR between 69.75 and 94.25 bpm; and 400 ms after the R-peak for an instantaneous HR higher than 94.25 bpm (Fittipaldi et al., 2020; García-Cordero et al., 2017; Yoris et al., 2017). A response was considered accurate if it fell within the defined temporal windows. The interoceptive accuracy index was calculated with the formula:

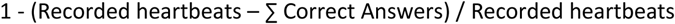

The interoceptive accuracy score range falls between 0 and 1, and higher scores indicate better interoceptive performance.

In the EC, we calculated the exteroceptive accuracy index by time-locking each tapped response with the corresponding time window for each presented heartbeat sound. A tapped response was considered correct if it fell between 0 to 750 ms from the recorded heartbeat (Fittipaldi et al., 2020; García-Cordero et al., 2017; Yoris et al., 2017). As for the interoceptive score, the exteroceptive accuracy index was calculated with:

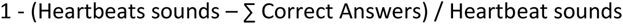

Mean latencies were also calculated as the average time between every correct tapped response and its corresponding recorded heartbeat (IC) and heartbeat sound (EC). Finally, both IC and EC tapped responses were assigned to “inhale” if the respective R-peak fell within the inhale onset/inhale offset time range and to “exhale” if the R-peak fell within the exhale onset/exhale offset time range. Using the same procedures described above, we calculated interoceptive and exteroceptive accuracy scores and mean latencies corresponding to the inhale and exhale phase of respiration (e.g., “interoceptive accuracy-inhale”, “interoceptive accuracy-exhale”, “exteroceptive accuracy-inhale”, “exteroceptive accuracy-exhale”, etc.). An overview of the experimental procedure and data analysis is presented in Figure 1. We tested for changes in task accuracy and mean latencies among respiratory phases (inhale vs. exhale) with a bootstrapped paired t-test (2000 permutations) within both the EC and the IC of the HBD task. Then, to investigate possible interaction effects of task by respiratory phase on task accuracy, we performed a two-way repeated-measures ANOVA on task accuracy with task condition (exteroceptive vs. interoceptive) and respiratory phase (inhale vs. exhale) as within-participants factors.

**Figure 1.**
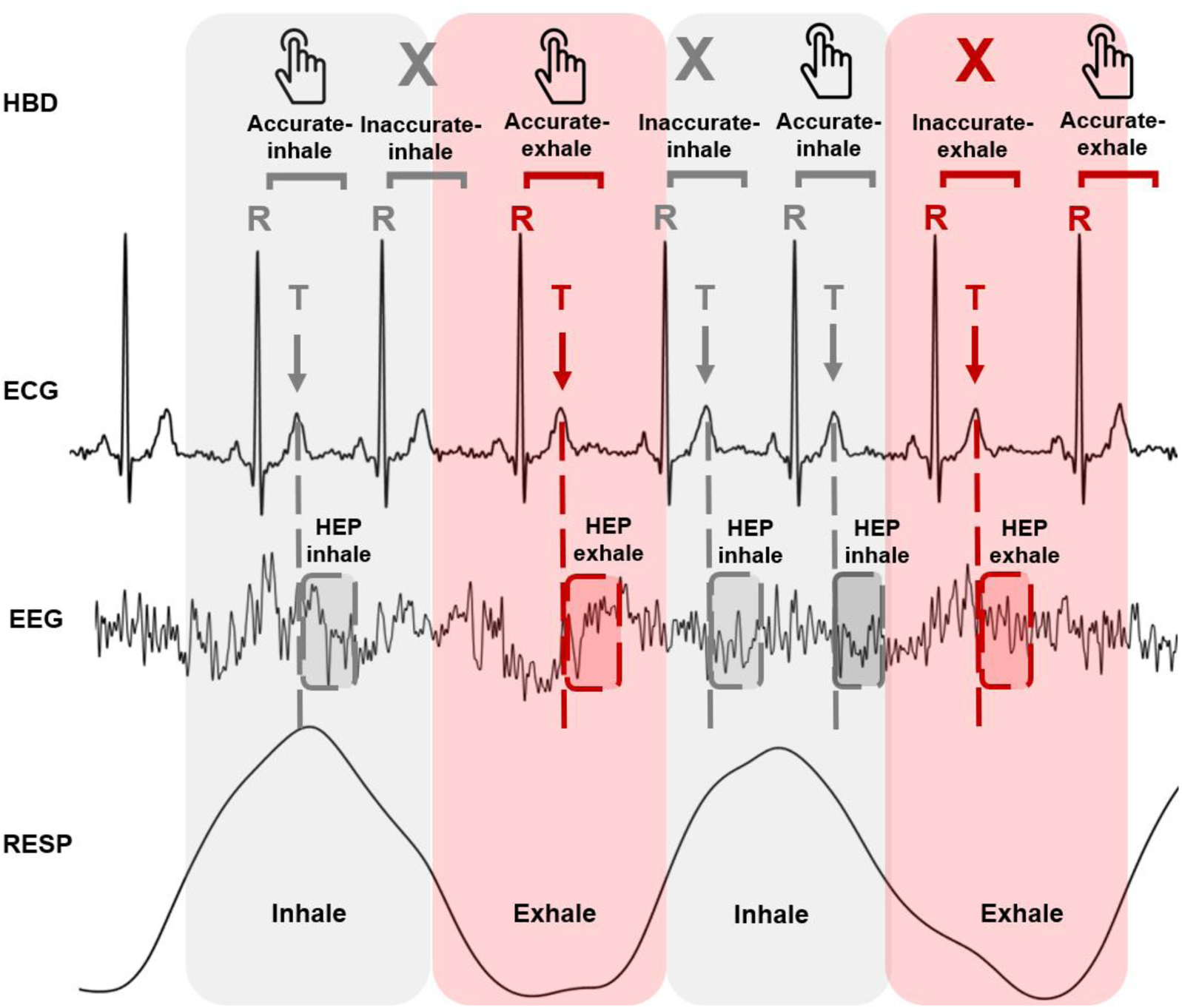
Schematic representation of data analysis. Grey areas represent inhale phases, pink areas represent exhale phases. The HBD row refers to the performance at the Heartbeat Detection task. “Hand” icons represent correct tapped responses. “X” icons represent inaccurate responses. The ECG row represents an exemplary trace of the electrocardiogram. Correct tapped responses were defined as the responses occurring within a given time window time-locked to the previous R-peak of the ECG signal. The width of the time window was individualized, based on the heart rate of each participant. The EEG (electroencephalogram) row represents an exemplary trace of one EEG channel. EEG data were epoched and time-locked to the T-peak (epoch length: −100 to 350 ms). Grey and red squares superimposed on the EEG trace represent the time window where HEP was analysed. We analysed HEP epochs occurring either during inhalation or exhalation, excluding those across respiratory phases. RESP row represents an exemplary trace of the respiratory activity.

### HEP amplitude statistical analysis

HEP differences between respiratory phases were statistically assessed as in Study 1. To investigate the interaction effect of task and respiratory phase on mean HEP amplitude, we performed a two-way repeated-measures ANOVA with mean HEP amplitude as dependent variable, and task condition (exteroceptive vs. interoceptive) and respiratory phase (inhale vs. exhale) as within-participants factors. In addition, to relate inter-individual differences in the effect of respiratory phase on the HEP to changes in interoceptive accuracy, we calculated the participants’ mean HEP difference (ΔHEP) between respiratory phases, by subtracting the mean HEP during inhale from the mean HEP during exhale in the significant time-window, averaged within the cluster of significant electrodes, as in Study 1. We next calculated participant’s interoceptive accuracy difference (Δaccuracy) between respiratory phases by subtracting the accuracy scores of the inhale phase (“interoceptive accuracy-inhale”) from the accuracy scores of the exhale phase (“interoceptive accuracy-exhale”). Then, based on previous evidence (Katkin et al., 1991; Mai et al., 2018; Montoya et al., 1993; Pollatos and Schandry, 2004; Schandry and Montoya, 1996), we tested the linear positive relationship between participants’ Δaccuracy and ΔHEP using Pearson’s correlation analysis.

### Additional cardiorespiratory analyses during the Heartbeat Detection Task

To investigate the role of cardiac and respiratory physiology on HEP activity and task accuracy, we performed the following additional analyses (Petzschner et al., 2019). Differences in the ECG amplitude between the IC and EC of the HBD task were first tested to exclude confounding cardiac effects. We compared the averaged ECG signal time-locked to the T-peak (epoch length: −100 to 350 ms) between the EC and IC using a t-test with FDR correction for all time points within the time window of observed significant HEP differences (Groppe et al., 2011). Then, differences in cardiorespiratory features among the EC and IC were assessed using bootstrapped paired t-test (2000 permutations). Within the IC, we also tested if the cardiac activity was different between respiratory phases (inhale vs. exhale), as in Study 1 (Groppe et al., 2011). Finally, we tested for any relationships between mean HEP differences (ΔHEP), interoceptive accuracy changes (Δaccuracy), and cardiorespiratory features of interest (Supplementary Material 1) across participants with a series of Pearson’s correlation analyses.

Throughout the manuscript, p-values (one for each feature) were adjusted for multiple testing using Benjamini and Yekutieli procedure (FDR, Benjamini and Yekutieli, 2001). FDR threshold was set at p = 0.05. All statistical analyses were performed in jamovi (v2.2.2; The jamovi project, 2021).

## Results

### Study 1 – Resting-state condition

#### Overview

We characterized respiratory phase-dependent HEP activity changes by comparing HEP activity between inhalation and exhalation during a 10-minutes resting-state condition. We used a comprehensive approach that included the entire EEG sensor space and epoch time window (Petzschner et al., 2019). The homogeneity of the inhale and the exhale phases in terms of number of registered heartbeats and analysed epochs was first verified by performing between-phases paired t-test comparisons (Supplementary Material 2).

### Heartbeat-evoked potentials activity among respiratory phases at rest

We determined whether a HEP occurred during inhale or exhale. We then performed a cluster-based permutation t-test to compare HEPs occurred over the whole scalp during the two respiratory phases, between 80 and 350 ms following the T-peak. In a time window ranging from 180 to 350 ms after the T-peak, HEP showed increased positivity, during exhalation, in a wide cluster of frontal, central, and parietal electrodes (FC1, FC2, FC3, Cz, C1, C2, C3, C4, CPz, CP1, CP2, CP3, CP4, Pz, P1, P3, P4, POz), peaking on CPz (cluster-based permutation t-test, 10000 permutations, t(38) = 2.0248, p_corrected_ = 0.0074) (Figure 2A-B). Source reconstruction analysis with sLORETA showed that HEP amplitude was significantly different between the two respiratory phases in two postero-central cortical areas, one on the left hemisphere (cluster-based permutation *t-*test, 1000 permutations, p_corrected_ = 0.002, cluster statistic (maxsum) = 13494, cluster size = 3904), while the other on the right hemisphere (cluster-based permutation *t-*test, 1000 permutations, p_corrected_ = 0.004, cluster statistic (maxsum) = 12089, cluster size = 3662). Such respiratory phase-dependent changes of HEP activity mapped onto areas of the Sensorimotor Network (bilateral post-central, paracentral, and pre-central gyrus) and the Default Mode Network (left inferior parietal lobule, and bilateral precuneus, cuneus, intraparietal sulcus, parieto-occipital cortex, and parietal superior lobule) (Figure 2C).

**Figure 2.**
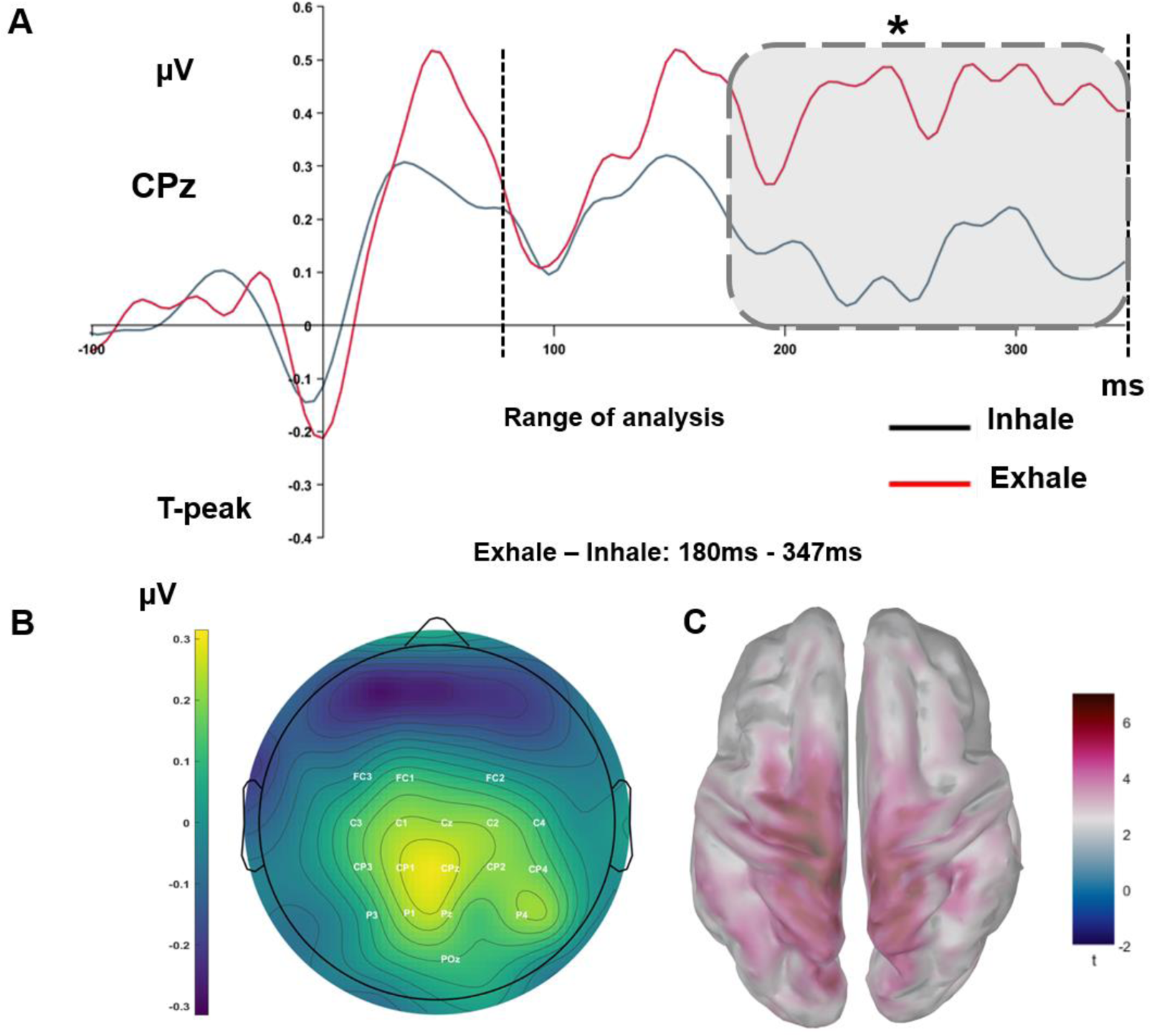
HEP activity changes among respiratory phases at rest. (A) Grand-average HEP waveforms at CPz. HEP waveform occurring at inhale is indicated in black, HEP waveform occurring at exhale is indicated in red. The dotted lines represent the temporal window of interest used for the statistical analysis (80-350 ms after the T-peak) of the HEP components. The grey rectangle marks the time window of significant differences (180–347 ms after the T-peak) (cluster-based permutation test). (B) Topographical scalp distribution showing the contrast between significant mean HEP differences (180–347 ms after the T-peak) between the exhale and inhale phases. (C) sLORETA source-reconstruction of significant HEP amplitude changes between the exhale and inhale phases in the Sensorimotor Network and the Default Mode Network (cluster-based permutation test).

### Additional cardiorespiratory analyses during resting-state

Heart physiology significantly differs between inhalation and exhalation (Draghici and Taylor, 2016; Riganello et al., 2018; Shaffer et al., 2014; Shaffer and Venner, 2013). Hence, possible confounding influence of CFA needs to be checked and ruled out in our study. Therefore, we performed additional analyses to support the assumption that observed differences in HEP amplitude between inhalation and exhalation were not driven by CFA modifications between phases, rather they reflected differences in heartbeat cortical processing (Petzschner et al., 2019). First, we tested for differences in ECG amplitude across the two phases with a repeated-measures two-tailed t-test at all time points within the time window of significant HEP differences, followed by FDR correction for multiple comparisons. We found statistically significant differences between inhale and exhale in ECG amplitude, characterized by increased ECG negativity in the exhale phase (paired t-test, t(38) = −4.674, p_FDR_ < 0.001) (Supplementary Figure 1A). This result suggests that the heartbeat itself was different during exhalations compared to inhalations. We, therefore, tested whether individual differences in ECG signal among respiratory phases (ΔECG) correlated with the effect of respiratory phase on HEP (ΔHEP) with a bootstrapped Pearson’s correlation (2000 permutations). There was no significant correlation between ΔECG and ΔHEP (Pearson’s correlation ΔECG, r = 0.222, uncorrected p = 0.175 (Supplementary Figure 1B), excluding a direct role of heartbeat physiology in explaining respiratory phase-dependent HEP modulations. We then estimated relevant cardiorespiratory features in our participants during the resting-state (Supplementary Table 1). Specifically, we measured HR, HF power, HRV total power, LF/HF ratio, as cardiac features, and breathing rate, inhale duration, exhale duration, I/E ratio, as respiratory features. Then, we performed a correlation analysis testing for any relationship between the observed mean HEP differences among respiratory phases (ΔHEP) and each cardiorespiratory feature of interest. Finally, we also checked possible associations between ΔHEP and, first, the number of registered heartbeats, second, the number of analysed epochs. All the correlation analyses resulted in null findings (Supplementary Material 3).

### Study 2 – Heartbeat Detection Task

#### Overview

We tested HEP activity changes across the respiratory cycle in participants performing the IC and the EC of the HBD task. During EC, participants heard a digital heartbeat sound and had to tap a button with their dominant hand in synchrony with it. During IC, they were asked to focus their attention on their own heart and tap a button at each heartbeat. As in Study 1, we verified the homogeneity of the inhale and the exhale phases in terms of number of registered heartbeats and analysed epochs by performing between-phases paired t-tests, both within the EC and the IC of the HBD task (Supplementary Material 4).

### Heartbeat-evoked potentials change across the respiratory cycle during IC

We first tested our hypothesis that HEP activity is modulated by respiratory phases during IC. As in Study 1, we first discriminated between HEP occurred during inhale and exhale. Then, we performed a cluster-based permutation t-test comparing HEPs occurring during the two respiratory phases, between 80 and 350 ms after the T-peak over the whole scalp. The results showed that during a time window ranging from 164 to 350 ms following the T-peak, HEP increased in positivity in a cluster of central and frontal electrodes (F2, F4, FC2, FC4, FC6, Cz, C1, C2, C4, C6, CPz, CP2, FT8), peaking on CPz and CP2. This result suggests that cortical processing of the heartbeat increases during exhalation phase, compared to inhalation phase, when participants focused their attention on heartbeats (cluster-based permutation *t-*test, 10000 permutations, t(18) = 2.1018, p_corrected_ = 0.0136) (Figure 3A-B, IC).

**Figure 3.**
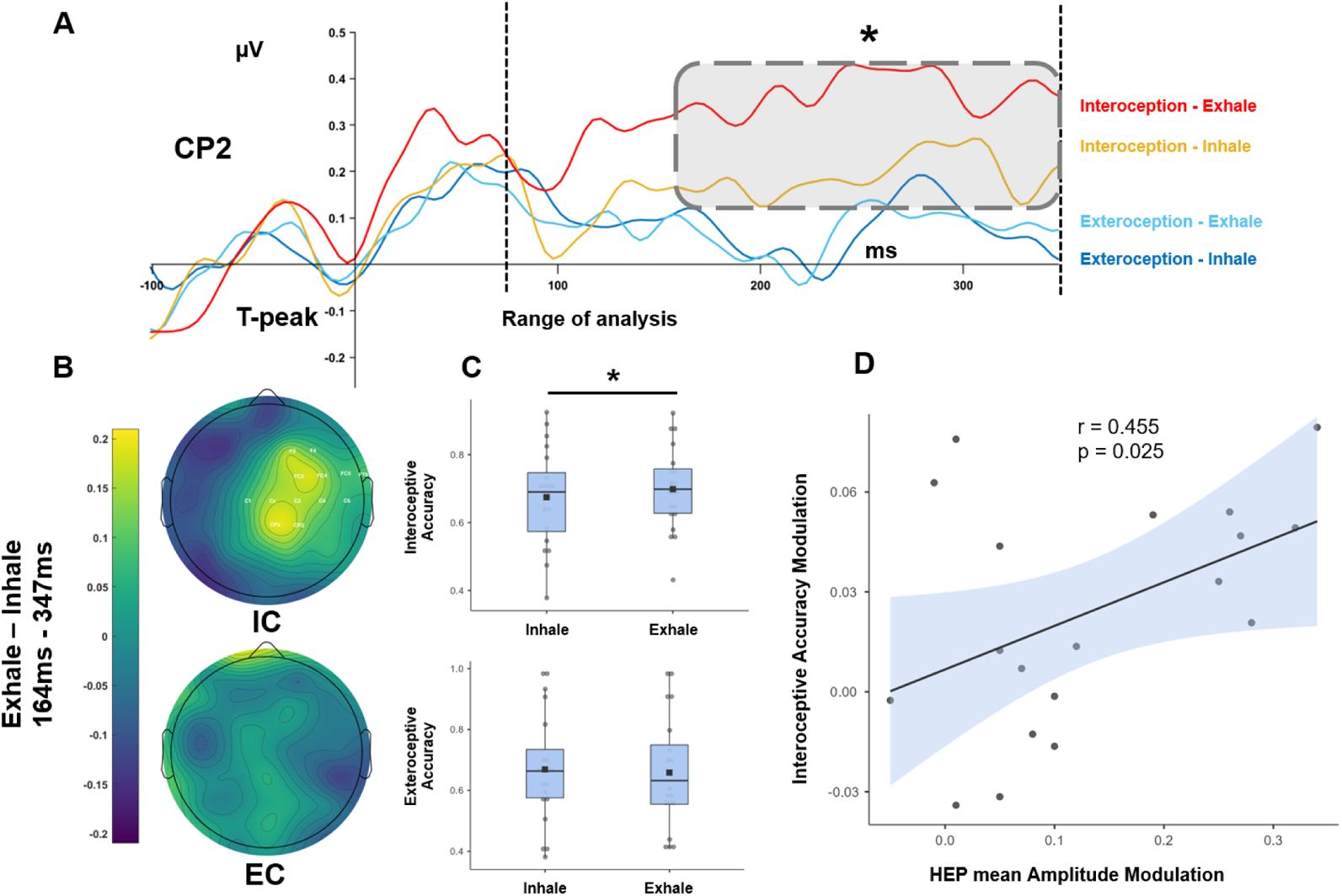
HEP activity and interoceptive accuracy change across the respiratory cycle during the HBD task. (A) Grand average HEP waveform at CP2. The time courses of the HEP are shown for “Interoception-Exhale” (red), “Interoception-Inhale” (orange), “Exteroception-Exhale” (light blue), and “Exteroception-Inhale” (dark blue). The dotted lines represent the temporal window of interest used for the statistical analysis (80-350 ms after the T-peak) of the HEP components. The grey rectangle marks the time window of significant differences (180–347 ms after the T-peak) (cluster-based permutation test). (B) Topographical scalp distribution (164–347 ms after the T-peak) shows the difference between the exhale and inhale phases for both the Interoceptive Condition (up) and the Exteroceptive Condition (down) of the HBD task. (C) Interoceptive (up) and Exteroceptive (down) accuracy scores of the HBD task for tapped responses plotted against the respiratory phases. * represents significant differences between interoceptive accuracy scores at inhale and exhale. (D) Scatter plot of the linear relationship (Pearson’s correlation, one tail) between the mean HEP amplitude changes (ΔHEP) and interoceptive accuracy changes (Δaccuracy) among respiratory phases. Mean HEP amplitude changes (ΔHEP) were computed by aggregating the following channels: F2, F4, FC2, FC4, FC6, Cz, C1, C2, C4, C6, CPz, CP2, FT8.

Second, we tested whether similar HEP modulations among respiratory phases are present also during the EC. To this aim, we compared HEP activity during exhalation and inhalation phases of respiration also in participants performing the EC. We adopted two different approaches: first, we compared inhale and exhale HEP over the whole scalp and in the whole time window (80-350 ms), as in IC; second, we focused on space regions of the main respiratory effects previously detected within the IC (i.e., F2, F4, FC2, FC4, FC6, Cz, C1, C2, C4, C6, CPz, CP2, FT8), in the significant time-window (164-350 ms). In both cases, cluster-based permutation t-tests revealed no differences in any cortical cluster (cluster-based permutation *t*-test, 10000 permutations, p_corrected_ > 0.05) (Figure 3A-B, EC). This null effect during EC suggests that respiratory phase-dependent HEP changes observed during IC are specifically associated to the direction of top-down attention: towards one’s own heartbeat (IC) rather than an external digital heartbeat (EC).

To investigate possible interaction effects of the two independent variables task and respiratory phase on mean HEP amplitude, we performed a two-way repeated-measures ANOVA with mean HEP amplitude as dependent variable, while task condition (exteroceptive vs. interoceptive) and respiratory phase (inhale vs. exhale) were the within-participants factors. The ANOVA did not show a main effect of task (F(1, 18) = 0.017, p = 0.899). Differently, the main effect of respiratory phase (F(1, 18) = 5.153, p = 0.036) was significant, as well as the task by respiratory phase interaction (F(1, 18) = 7.517, p = 0.013). This result suggests that respiratory phase-dependent modulation of HEP is significantly stronger when top-down attention is directed towards the heart than towards the external heartbeat-like sounds.

### Interoceptive accuracy changes across the respiratory cycle

We hypothesized that, along with HEP activity, interoceptive accuracy is modulated by respiratory phases. Therefore, we tested if accuracy of tapped responses given during IC in the exhale phase of respiration was different from accuracy of tapped responses given in the inhale phase of the same task condition. First, we calculate participants’ interoceptive and exteroceptive accuracy and mean latency (Supplementary Material 5). Then, we separately analysed interoceptive and exteroceptive accuracy scores and mean latencies associated to the inhale and exhale phases of respiration. Mean interoceptive accuracy was 69.84 ± 12.56 % [mean ± SD] for the exhale phase, and 67.45 ± 14.71 % [mean ± SD] for the inhale phase. In line with HEP findings, a bootstrapped paired t-test (2000 permutations) revealed significant changes in interoceptive accuracy among respiratory phases, indicating increased detection of the heartbeat sensations during exhalations (paired *t*-test, t(19) = 3.14, p = 0.005) (Figure 3C - IC). Mean latency for accurate responses was the same among respiratory phases, being 335.11 ± 36.77 ms [mean ± SD] for the inhale phase, and 336.68 ± 41.20 ms [mean ± SD] for the exhale phase (paired *t-*test, t(19) = 0.223, p = 0.86).

To test the hypothesis that differences in accuracy among respiratory phases were specific to interoception, as for IC, we calculated exteroceptive accuracy in the EC during exhale and compared it to accuracy in the inhale phase of respiration. Mean exteroceptive accuracy was 64.10 ± 16.98 % [mean ± SD] for the exhale phase, and 65.23 ± 16.94 % [mean ± SD] for the inhale phase. We did not find any significant modulation of exteroceptive accuracy due to respiratory phases during EC (t(19) = 1.35, p = 0.19) (Figure 3C - EC). This result suggests that respiration does not affect the exteroceptive counterpart of the HBD task. As for IC, mean latency for accurate responses did not change significantly among respiratory phases, being 194.81 ± 66.32 ms [mean ± SD] for the inhale phase, and 200.64 ± 74.27 ms [mean ± SD] for the exhale phase (t(19) = −0.682, p = 0.50).

Next, we performed a two-way repeated-measures ANOVA with task accuracy as dependent variable, while task condition (exteroceptive vs. interoceptive) and respiratory phase (inhale vs. exhale) were the within-participants factors. The ANOVA showed no significant main effect of respiratory phase (F(1, 19) = 2,263, p = 0.150), nor of task condition (F(1, 19) = 0.583, p = 0.455). However, the interaction task by respiratory phase was significant (F(1, 19) = 6.464, p = 0.020). Crucially, this result confirms that, as for HEP activity, respiratory phase-dependent modulations of accuracy are specific to the interoceptive task.

Most interestingly, in agreement with existing evidence that HEP activity is positively associated to accuracy in heartbeat detection (Canales-Johnson et al., 2015; Marshall et al., 2017; Pollatos et al., 2005), and according to our hypothesis that such positive association may reflect also consistent modulation of HEP and interoceptive accuracy by respiratory phases, we found significant correlation between interoceptive accuracy changes (Δaccuracy) and changes of HEP mean amplitude (ΔHEP; r = 0.455, p = 0.025, one tail): the stronger the respiratory phase-dependent modulation of HEP activity, the stronger the respiratory-dependent modulation of interoceptive accuracy in individuals performing IC (Figure 3D).

### Additional cardiorespiratory analyses during the Heartbeat Detection Task

In order to control if our results were determined by differences in heart activity between IC and EC, we first tested for differences in ECG amplitude across conditions with a repeated-measures two-tailed t-tests at all time points within the time window of significant HEP differences (80-350 ms) followed by FDR correction for multiple comparisons. We did not find differences between IC and EC in the ECG signals (paired *t*-test, p_FDR_ = 0.318) (Supplementary Figure 1D). Then, within the IC, we tested if the CFA equally impacted the HEP among respiratory phases (inhale vs. exhale) in the time window of significant HEP differences (IC: 164 - 350 ms). We observed no differences in cardiac activity between inhale and exhale phases (paired *t*-test, p_FDR_ = 0.28) (Supplementary Figure 1C). We then compared participants’ cardiorespiratory features during the EC and the IC of the HBD task (Supplementary Table 2) with a bootstrapped paired t-test (2000 permutations) to rule out the possibility that any difference found between the two tasks at neural and behavioural levels could be merely explained by physiological contributions. Participants had lower respiratory rate during the IC, compared to EC (breathing rate: t(19) = −3.93, uncorrected p = 0.001, p_FDR_ = 0.022; inhale duration: t(19) = 3.51, uncorrected p = 0.002, p_FDR_ = 0.022; exhale duration: t(19) = 3.03, uncorrected p = 0.006, p_FDR_ = 0.043). Whether these changes were due to the different arousing nature of the two conditions, or to spontaneous adjustments of respiration to improve heartbeat perception cannot be determined based on present data. However, differences in breathing rate across IC and EC were not associated to observed respiratory phase-dependent modulations of HEP during IC (Supplementary Material 6). No other cardiorespiratory feature (HR, HF power, HRV total power, LF/HF ratio, I/E ratio) significantly differed between IC and EC (Supplementary Table 2). Finally, likewise in Study 1, we performed a series of correlation analyses during IC testing for relationships between i) mean HEP differences (ΔHEP) and cardiorespiratory features (Supplementary Material 7); and ii) interoceptive accuracy changes (Δaccuracy) and cardiorespiratory features (Supplementary Material 8). All the above-mentioned analyses did not yield statistically significant results, suggesting that observed differences in HEP activity and interoceptive accuracy among respiratory phases during IC cannot be simply explained by differences at the level of cardiac and respiratory physiology.

## Discussion

The fundamental influence of respiration on brain activity and cognitive functions in humans has been increasingly recognized in the last few years (Heck et al., 2017; Kluger and Gross, 2021; Varga and Heck, 2017). However, its potential role in fine-tuning brain-heart interactions has gone mostly unstudied within the interoception research field so far. In the present work, we performed two studies with the general aim of investigating the role of spontaneous respiration in shaping the cortical processing of cardiac-related information. In both studies, we focused on HEP activity modulations, an EEG event-related potential time-locked to the ECG signal, commonly regarded as an objective electrophysiological index of brain-heart interactions (Coll et al., 2021; Park and Blanke, 2019a).

### Neuro-cardio-respiratory interactions at rest

Our first specific aim was to investigate the interplay between the cardiac, respiratory and brain activity in a resting-state condition, that is, while participants were let mind-wander and spontaneously breath. Hence, in Study 1, we recruited healthy volunteers and computed HEP levels separately for heartbeats occurred during the inhalation and the exhalation phases of the respiratory cycle. We found higher HEP amplitude during exhalation compared to inhalation, indicating increased brain-heart interactions and improved cortical processing of the heartbeats. HEP levels significantly increased in a time window ranging from 180 to 350 ms after the T-peak over frontal, central, and parietal electrodes, and peaked on CPz. Source-level respiratory effects on HEP were localized in cortical regions overlapping the Sensorimotor Network and the Default Mode Network. Specifically, they included the left inferior parietal lobule, the bilateral post-central, paracentral, and pre-central gyri, the precuneus, cuneus, intraparietal sulcus, superior parietal lobule, and parieto-occipital cortex. These results are consistent with previous evidence showing higher HEP activity over medial posterior areas associated to self-related processes during mind-wandering (Babo-Rebelo et al., 2016).

A first, low-level interpretation of the present results may refer to the strong coupling between the respiratory cycle and the baroreflex. This well-known interaction is commonly referred to as RSA, a form of cardiorespiratory synchronization resulting in cyclic HR increases during inhalations and subsequent decreases during exhalations (Brecher and Hubay, 1955; Eckberg et al., 1985). RSA is mechanically induced by inhalation-dependent decrease of pleural pressure driven by chest expansion, which facilitates venous return to the right heart. This causes a reduction of the left ventricular stroke volume, decreasing aortic blood pressure, which in turn is sensed by aortic baroreceptors. Lower baroreceptor stimulation during inhalation then triggers the baroreflex: to stabilize cardiac output over the short term, the HR temporarily increases (Bainbridge, 1915; Magder, 2018). However, this also means that when HR increases, heartbeats are inherently weaker (Draghici and Taylor, 2016; Riganello et al., 2018; Shaffer et al., 2014; Shaffer and Venner, 2013). Regarding the present study, it is possible that being inhalations associated to increases in HR, they are also inextricably linked to weaker (or less salient) heartbeats, eventually resulting in the observed reduction of the HEP during inhalations compared to exhalations. Accordingly, a recent behavioural study found that the performance to a modified Heartbeat Tracking task was modulated by fluctuations in the frequency of heartbeats: participant’s interoceptive accuracy was higher when HR was lower (Larsson et al., 2021). To further explore this hypothesis, we performed a series of control analyses on both the ECG signal and cardiorespiratory features. On the one hand, supporting the interpretation that increased HEP levels may result from stronger heartbeats during exhalation, we found increased amplitude of the ECG signal, time-locked to the T-peak, in the exhalation compared to the inhalation phase. On the other hand, differences in the ECG signal did not correlate with mean HEP changes among respiratory phases, likely suggesting that the two phenomena are likely independent from each other. Additionally, respiratory phase-related HEP changes did not correlate with any other cardiorespiratory feature possibly interfering with it, such as HR, HRV total power, and HF power (i.e., an indicator of RSA; Lewis et al., 2012). Hence, we suggest that heart physiology plays a minor role in the observed respiratory phase-dependent modulations of HEP activity.

Another possible interpretation of our results is that afferent interoceptive information travelling from the heart to the cerebral cortex could be modulated by simultaneous sensorimotor signals generated by pulmonary afferents as well as by the active movement of the chest (see Baumert et al., 2015 for a similar interpretation). Indeed, heartbeat and respiratory interoceptive signals follow similar pathways, as both aortic baroreceptors and rapidly-adapting pulmonary stretch receptors send information about the state of the cardiac and respiratory systems through the vagal nerve and the nucleus of the solitary tract (Park and Blanke, 2019a). It is worth highlighting here that respiratory sensorimotor and interoceptive information is sent upstream primarily during inhalations (Noble and Hochman, 2019; Streeter et al., 2012). Therefore, in this specific respiratory phase, weaker cardiac baroreceptor-mediated information may compete with, and likely lose against, stronger respiratory-mediated information. This would result in increased respiratory sensorimotor interference over cardiac signals, and, possibly, a reduced encoding of cardiac information. Supporting this hypothesis, we found that source-level HEP reduced activity during inhalations, compared to exhalations, was localized over brain regions that have been classically associated to respiratory phase-dependent fMRI modulations, that is, the Sensorimotor Network (Bijsterbosch et al., 2017; Birn et al., 2006; Glasser et al., 2018; Power et al., 2020).

A third intriguing explanation of our results relies on the predictive coding model of interoceptive perception (Barrett and Simmons, 2015; Seth, 2013; Seth and Friston, 2016). In brief, recent interoceptive inference models (Allen et al., 2019; Allen et al., 2021) posit that cardiorespiratory interoception is able to shape the neural gain (i.e., the balance of neural excitation vs. inhibition) across several brain regions by modulating the computational precision (i.e., the inverse of noise) of perceptual, cognitive, and emotional processes. More in detail, periodic physiological changes related to rhythmic cardiorespiratory oscillations are computed by the brain as stable predictions and are centrally suppressed via a sensory attenuation process in order to minimize their interference. Accordingly, recent studies (Al et al., 2020, 2021; Grund et al., 2021) on heart functions found that the systolic phase of the cardiac cycle (i.e., when baroreceptor activity is at a maximum) was related to simultaneous attenuation of somatosensory perception. This effect was explained by the functional cortical overlapping between somatosensory perception and cardiac interoception, at the level of the primary somatosensory cortices: in these regions, the same sensory attenuation process that minimized systole-related oscillations also reduced somatosensory processing (Al et al., 2020, 2021). Similarly, we propose that the brain receives recursive and predictable interoceptive signals during inhalations forming stable predictions about each respiratory cycle, and consequently suppressing the physiological signals associated to it within the Sensorimotor Network (Birn, 2012). Since sensorimotor areas are also involved in heartbeat-related information processing (Park and Blanke, 2019a), cardiac-related sensations of heartbeats occurring during inhalations could be suppressed together with those related to respiration. This would explain the observed decrease of HEP activity during inhalations within this network.

### Neuro-cardio-respiratory interactions during interoceptive and exteroceptive attentional tasks

Our second specific aim was to explore if the above-described neuro-cardio-respiratory interactions are further modulated by endogenous attention towards interoceptive vs. exteroceptive signals and if this modulation affects individual interoceptive vs. exteroceptive accuracy at behavioural level. In Study 2 we simultaneously recorded EEG, ECG, and respiratory activity in healthy volunteers during the IC and the EC of the HBD task (Fittipaldi et al., 2020; García-Cordero et al., 2017; Yoris et al., 2017). We observed significant respiratory phase-dependent modulation of HEP activity exclusively during IC. As in Study 1, HEP activity increased during exhalations in a time window ranging from 164 to 350 ms after the T-peak. This effect was found over a cluster of central and frontal electrodes, peaking at CPz and CP2. During EC, we observed no effects of the respiratory phase on HEP activity, suggesting that when attention is focused on the external environment, heart-brain interactions are no longer modulated by respiratory phases. Similarly, at the behavioural level, we found that the exhalation phase of respiration was beneficial exclusively for interoceptive accuracy, while exteroceptive accuracy was not modulated by respiratory phases. Direct comparisons of respiratory phase-induced modulations of HEP (ΔHEP) and accuracy (Δaccuracy) between the interoceptive and exteroceptive conditions showed higher ΔHEP and higher Δaccuracy during the IC, compared to the EC. Interestingly, these modulations were positively correlated. As in Study 1, control analyses on both the ECG signal and relevant cardiorespiratory features ruled out the possibility that modulations of HEP activity and interoceptive accuracy could be explained by confounding influences of the CFA, or merely at the level of cardiorespiratory physiology. To summarize, Study 2 confirms the findings of Study 1 and further extends them by showing that cortical processing and perception of heartbeats are optimized at exhalation when attention is directed towards interoceptive signals (see Herrero et al., 2018 for similar results on interoception and EEG-breath coherence).

To better understand our results, we can interpret the role of attention towards the heartbeat during IC and towards the environment during EC within the framework of the predictive coding model. According to this model, top-down attention increases the precision of what is relevant for the organism in a specific moment, by modulating neuronal gain that represents the target objects at the expense of others (Feldman and Friston, 2010; Smout et al., 2019; Boyadzhieva and Kayhan, 2021). Then, during IC, the brain may assign the highest priority to heartbeat signals; on the opposite, during EC, the brain may assign the highest priority to the heard sounds. Notably, however, during both IC and EC the brain also receives recursive, predictable, and strong respiratory-related noise during inhalations. Hence, the question arises of why we observed different respiratory phase-dependent modulations of HEP activity between the IC (higher ΔHEP) and the EC (lower ΔHEP). According to the above-described interoceptive inference models (Allen et al., 2019; Allen et al., 2021), interoceptive prediction is not an all-or-none phenomenon, but a highly context-sensitive process determining the precision of incoming sensations based on the ongoing task-oriented cognition. Therefore, during the IC, to be able to correctly perceive and process heartbeat sensations across the respiratory cycle, the brain may adaptively increase the precision of interoceptive cardiac input specifically during exhalations, that is, when inhalation-related physiological noise is absent. This is not necessary during the EC, because auditory signal processing at the level of auditory areas is much less interfered by cardiac and respiratory signal processing. Hence, interoceptive optimization of the heartbeat signal processing across the respiratory cycle may occur during the IC, reflected by higher HEP increases during exhalations (higher ΔHEP), but not during the EC (lower ΔHEP). Notably, this interpretation is further supported by the linear correlation between respiratory phase-induced modulation of both HEP and interoceptive accuracy, with higher ΔHEP (i.e., interoceptive optimization) associated to higher increase of cardiac interoceptive accuracy (Δaccuracy). This clearly suggests that the ΔHEP index herein reported for the first time reflects a degree of optimization of interoceptive processing that is also relevant for interoceptive perception.

### Conclusions and limitations of the study

Overall, the present findings reveal a so far unnoticed influence of respiration on cardiac interoception: when contextualized within a breathing organism, cardiac interoception is highly interconnected with the respiratory cycle, in addition to task-oriented cognition. Accordingly, we propose that the respiratory phase-dependent HEP modulation (ΔHEP) could represent a physiological index of cardiac interoceptive optimization, with behavioural implications, above and beyond HEP activity alone. In general, we underline the importance of investigating the synergic interplay between different visceral signals, which have been classically studied in isolation within the field of interoception research (Garfinkel et al., 2016). In particular, building on the present research, future studies should investigate neuro-cardio-respiratory interactions during different interoceptive and exteroceptive tasks. In fact, a limitation of the present study is that the occurrence of motor tapped responses during the HBD task made the comparison between ΔHEP during task vs. rest inappropriate, due to motor-related confounding activity intrinsic to the HBD task. To overcome this limitation, future studies could, for instance, adopt the Schandry task (Schandry, 1981), a non-motor-based cardiac interoceptive task, making direct comparisons of ΔHEP during rest and task possible. It could be also tested if voluntary breath-control, in particular the slowing down of the breathing frequency or the specific increase of the exhalation duration (i.e., the decrease of the I/E ratio), may modulate neurophysiological and behavioural signatures of cardiac interoception (MacKinnon et al., 2013). This would be particularly relevant to better understand the relationships between interoception and mental health, because breath-control is a highly recommended practice in a whole range of clinical applications and mind-body interventions (Farb et al., 2015; Paulus, 2013; Weng et al., 2021; Zaccaro et al., 2018, 2022). Finally, the link between ΔHEP and interoception could be tested in different clinical or sub-clinical populations with known altered or reduced interoceptive sensitivity, such as generalized anxiety disorder, major depression, and schizophrenia (Ardizzi et al., 2016; Bonaz et al., 2021).

To conclude, in line with the active sensing interpretation of respiratory activity, we speculate on the existence of a breathing-related attentional “switch” between interoception and exteroception. It is already known that inhalation leads to adaptive modulation of neuronal gain in order to facilitate reception and elaboration of sensory information from the external environment (Grund et al., 2021; Huijbers et al., 2014; Kluger et al., 2021; Perl et al., 2019; Zelano et al., 2016). As inhalation itself improves exteroception-related cognitive functions, exhalation may reflect a more general attentional shift towards the internal bodily states and the self, hence modulating interoceptive perception as well as self-consciousness (Molle and Coste, 2022; Park et al., 2018; Park and Blanke, 2019b). That is, spontaneous breathing may continuously tune the brain to switch from extrinsically (external) oriented processing during inhalation, to intrinsically (internal) oriented functions during exhalation (Golland et al., 2007).

## Acknowledgements

This study was supported by the “Departments of Excellence 2018-2022” initiative of the Italian Ministry of Education, University and Research for the Department of Neuroscience, Imaging and Clinical Sciences (DNISC) of the University of Chieti-Pescara, and by the “Search for Excellence” program, University of Chieti-Pescara. We thank Valentina Torzolini for her support in data acquisition.

