## Supplementary Material for "Brain-heart interactions are optimized across the respiratory cycle via interoceptive attention"

### Supplementary Material 1. Cardiorespiratory features of interest

**Breathing rate** (breaths/min): the average number of breathing cycles in a minute

**Inhale duration** (sec): the average duration of the inhalation phases of respiration

**Exhale duration** (sec): the average duration of the exhalation phases of respiration

**Inhalation/Exhalation** (I/E) ratio: the ratio between the average inhale duration and the average exhale duration

**Heart Rate** (HR) (beats/min): the average number of heartbeats in a minute

**Heart Rate Variability** (HRV) total power ($ms^2$/Hz): the sum of the power values in HF (0.15-0.4 Hz), LF (0.04-0.15 Hz), and VLF (0-0.04 Hz) frequency bands. It estimates the degree of HRV in the frequency domain

**High Frequency** logarithmic power (HFlog) (log $ms^2$/Hz): log-transformed power of HF band (0.15-0.4 Hz). It is a marker of parasympathetic activation, and it is a main indicator of Respiratory Sinus Arrhythmia (RSA, i.e., respiratory cycle influences on HR; Lewis et al., 2012)

**Low Frequency/High Frequency ratio** (LF/HF): the ratio between LF and HF band powers. It is an estimated of sympatho-vagal balance. Its decrease is a marker of parasympathetic activity prevalence

### Supplementary Material 2. Testing the homogeneity of the inhale and the exhale phases

In total, we registered 19571 heartbeats, 9845 heartbeats (252 ± 78 [mean ± SD]) during inhalation, and 9726 (249 ± 85 [mean ± SD]) during exhalation. After artifact correction, we retained 4525 breathing cycles (113 ± 44.63 [mean ± SD]), and we analysed a total of 14415 HEP epochs, divided into 7061 epochs for the inhale phase (181 ± 56.8 [mean ± SD]), and 7354 epochs for the exhale phase (188 ± 69.36 [mean ± SD]). Both the number of detected heartbeats, and the number of analysed HEP epochs did not differ between exhalation and inhalation phases (paired *t*-test detected heartbeats, t(39) = 0.588, p = 0.56; paired *t*-test HEP epochs, t(38) = -1.483, p = 0.146), supporting the homogeneity of the inhale and the exhale phases.

**Supplementary Material 3. Correlations with cardiorespiratory features and with registered heartbeats/epochs**

We tested for significant correlation between subject's HEP changes (ΔHEP) and cardiorespiratory features of interest with a bootstrapped Pearson's correlation (2000 permutations). There was no significant correlation between subject's HEP changes (ΔHEP) and any cardiac features (Pearson's correlation HRV total power, r = -0.232, uncorrected p = 0.149; Pearson's correlation HFlog, r = -0.017, uncorrected p = 0.917; Pearson's correlation LF/HF, r = -0.173, uncorrected p = 0.286), with mean inhale duration (Pearson's correlation, r = -0.309, uncorrected p = 0.052), and with I/E ratio (Pearson's correlation, r = 0.199, uncorrected p = 0.219). We found a trend showing a mean HEP changes positive correlation with breathing rate and a negative correlation with mean exhale duration, which did not survive to FDR correction (Pearson's correlation breathing rate, r = 0.401, uncorrected p = 0.01, $p_{FDR}$ = 0.196; Pearson's correlation mean exhale duration, r = -0.371, uncorrected p = 0.018, $p_{FDR}$ = 0.196).

We also tested for relationships between ΔHEP changes and both the number of registered heartbeats and the number of retained artifact-free HEP epochs in any respiratory phase, as well as with their differences between inhale and exhale, finding null results (Pearson's correlation registered heartbeats inhale r = -0.153, uncorrected p = 0.353; Pearson's correlation registered heartbeats exhale r = -0.182, uncorrected p = 0.268; Pearson's correlation number of HEP epochs inhale r = -0.255, uncorrected p = 0.116; Pearson's correlation number of HEP epochs exhale r = -

0.254, uncorrected p = 0.119; Pearson's correlation Δregistered heartbeats r = 0.107, uncorrected p = 0.517; Pearson's correlation ΔHEP epochs r = 0.097, uncorrected p = 0.556).

**Supplementary Material 4. Testing the homogeneity of the inhale and the exhale phases for IC and EC**

We analysed a total of 6178 artifact-free respiratory cycles: 2640 breaths in the IC (132 ± 48.56 [mean ± SD]), and 3538 in the EC (177 ± 43.01 [mean ± SD]). In the IC, we registered a total of 10742 heartbeats, 5387 heartbeats (283 ± 82 [mean ± SD]) during inhalation, and 5355 (281 ± 82 [mean ± SD]) during exhalation. Of these, we retained for the analysis 7927 artifact-free HEP epochs: 3907 for the inhale phase (205 ± 61.17 [mean ± SD]), and 4020 for the exhale phase (211 ± 62.97 [mean ± SD]). In the EC, we registered a total of 12018 heartbeats, 6156 heartbeats (324 ± 76 [mean ± SD]) during inhalation, and 5862 (308 ± 69 [mean ± SD]) heartbeats during exhalation. After artifact rejection, we analysed 8252 HEP epochs, 4078 for the inhale phase (214 ± 50 [mean ± SD]), and 4174 for the exhale phase (219 ± 49 [mean ± SD]). Neither the total number of detected heartbeats, nor the number of retained HEP epochs differed in the contrast between IC and EC (paired t-test detected heartbeat, t(19) = -1.259, p = 0.224; paired t-test HEP epochs, t(18) = -0.459, p = 0.651). Both within the IC and the EC, the total number of detected heartbeats, and the number of retained HEP epochs did not differ between inhale and exhale phases (paired t-test detected heartbeat IC, t(19) = 0.181, p = 0.858; paired t-test HEP epochs IC, t(18) = -0.746, p = 0.465; paired t-test detected heartbeat EC, t(19) = 1.528, p = 0.144; paired t-test HEP epochs EC, t(18) = -0.448, p = 0.66), supporting the homogeneity of the inhale and the exhale phases for the IC and EC of the HBD task.

**Supplementary Material 5. Interoceptive and exteroceptive accuracy and mean latencies**

On average, participants' interoceptive accuracy was 68 ± 13% [mean ± SD], while mean exteroceptive accuracy was 66 ± 18% [mean ± SD]. Mean latency for accurate tapped responses was 334.95 ± 34.97 ms [mean ± SD] for the IC, and 196.09 ± 68.86 ms [mean ± SD] for the EC. Mean accuracy did not differ between IC and EC (paired t-test, t(19) = 0.365, p = 0.719), while mean latency for accurate responses was higher in IC than EC (paired t-test, t(19) = 8.911, p = 0.000).

**Supplementary Material 6. Correlations with cardiorespiratory features differences between EC and IC**

We tested if observed changes in breathing rate across the IC and EC were correlated with HEP modulations among respiratory phases during IC. There was no significant correlation between subject's HEP changes (ΔHEP) and breathing rate changes (Pearson's correlation breathing rate changes, r = 0.059, uncorrected p = 0.806; Pearson's correlation inhale duration changes, r = 0.241, uncorrected p = 0.307; Pearson's correlation exhale duration changes, r = 0.108, uncorrected p = 0.651).

**Supplementary Material 7. HEP differences and cardiorespiratory features**

We tested if changes in HEP activity among respiratory phases in the IC were correlated with individual differences in cardiorespiratory features of interest. There was no significant correlation between subject's HEP changes (ΔHEP) and any respiratory features (Pearson's correlation breathing rate, r = -0.345, uncorrected p = 0.148; Pearson's correlation inhale duration, r = 0.391, uncorrected p = 0.098; Pearson's correlation exhale duration, r = 0.344, uncorrected p = 0.149; Pearson's correlation I/E ratio, r = 0.054, uncorrected p = 0.826). We found a trend showing that mean HEP changes negatively correlated with HR and positively correlated with HRV total power, which did not survive to FDR correction (Pearson's correlation HR, r = -0.569, uncorrected p = 0.011,

$p_{FDR}$ = 0.12; Pearson's correlation HRV total power, r = 0.566, uncorrected p = 0.011, $p_{FDR}$ = 0.12). There was no correlation between ΔHEP, HFlog power, and LF/HF (Pearson's correlation HFlog, r = 0.298, uncorrected p = 0.215; Pearson's correlation LF/HF, r = 0.319, uncorrected p = 0.183).

**Supplementary Material 8. Interoceptive accuracy changes and cardiorespiratory features**

We checked whether changes in interoceptive accuracy during respiratory phases (Δaccuracy) in IC were correlated with individual differences in cardiorespiratory features of interest. We did not find significant correlations between subject's interoceptive accuracy changes and any respiratory features (Pearson's correlation breathing rate, r = -0.130, p = 0.586; Pearson's correlation inhale duration, r = 0.244, p = 0.301; Pearson's correlation exhale duration, r = 0.130, p = 0.584; Pearson's correlation I/E raio, r = 0.154, p = 0.517), as well as with mean HR (Pearson's correlation, r = 0.118, p = 0.621), and LF/HF (Pearson's correlation, r = 0.025, p = 0.916). We found a trend of positive correlations between Δaccuracy, HRV total power, and HFlog power, which did not survive to FDR correction (Pearson's correlation HRV total power, r = 0.456, uncorrected p = 0.043, $p_{FDR}$ = 0.467; Pearson's correlation HFlog power, r = 0.514, uncorrected p = 0.021, $p_{FDR}$ = 0.457).

**Supplementary Table 1. Cardiorespiratory features during resting-state**

| | Resting-state | |
|---|---|---|
| | **Mean** | **SD** |
| **Respiratory rate** (bpm) | 15.60 | 4.52 |
| **Average inhale duration** (sec) | 1.97 | 0.76 |
| **Average exhale duration** (sec) | 2.30 | 0.97 |
| **I/E ratio** | 0.87 | 0.14 |
| **Heart rate** (bpm) | 72.73 | 9.83 |
| **Heart rate variability** ($ms^2$/Hz) | 2341.36 | 2423.59 |
| **HF** (Log ms2/Hz) | 6.58 | 0.96 |
| **LF/HF** | 2.06 | 2.91 |

Abbreviations: I/E-Inhalation/Exhalation, HF-High Frequency, LF-Low Frequency, SD-Standard Deviation

**Supplementary Table 2. Cardiorespiratory features during the EC and the IC of the HBD task**

| | **Interoceptive Condition** | | **Exteroceptive Condition** | | **Interoceptive vs. Exteroceptive** | | |
|---|---|---|---|---|---|---|---|
| | **Mean** | **SD** | **Mean** | **SD** | **t** | **uncorrected p** | **$p_{FDR}$** |
| **Respiratory rate** (bpm) | 16.7506 | 4.1606 | 20.3224 | 3.7080 | 3.93 | 0.001 | **0.022** |
| **Average inhale duration** (sec) | 1.8432 | 0.5946 | 1.4817 | 0.3314 | -3.51 | 0.002 | **0.022** |
| **Average exhale duration** (sec) | 2.0433 | 0.7319 | 1.6027 | 0.4671 | -3.03 | 0.006 | **0.044** |
| **I/E ratio** | 0.9175 | 0.1175 | 0.9434 | 0.1359 | 1.06 | 0.301 | |
| **Heart rate** (bpm) | 70.6974 | 11.0702 | 70.9263 | 12.6210 | 0.12 | 0.907 | |
| **Heart rate variability** ($ms^2$/Hz) | 2060.51 | 1975.228 | 1302.436 | 895.154 | -2.81 | 0.011 | 0.06 |
| **HF** (Log $ms^2$/Hz) | 6.4459 | 0.7957 | 6.2873 | 0.8760 | -1.37 | 0.187 | |
| **LF/HF** | 1.7142 | 1.6785 | 1.1474 | 1.2445 | -2.02 | 0.056 | |

Abbreviations: HF-High Frequency, LF-Low Frequency, SD-Standard Deviation, FDR-False Discovery Rate

**Supplementary Figure 1.**

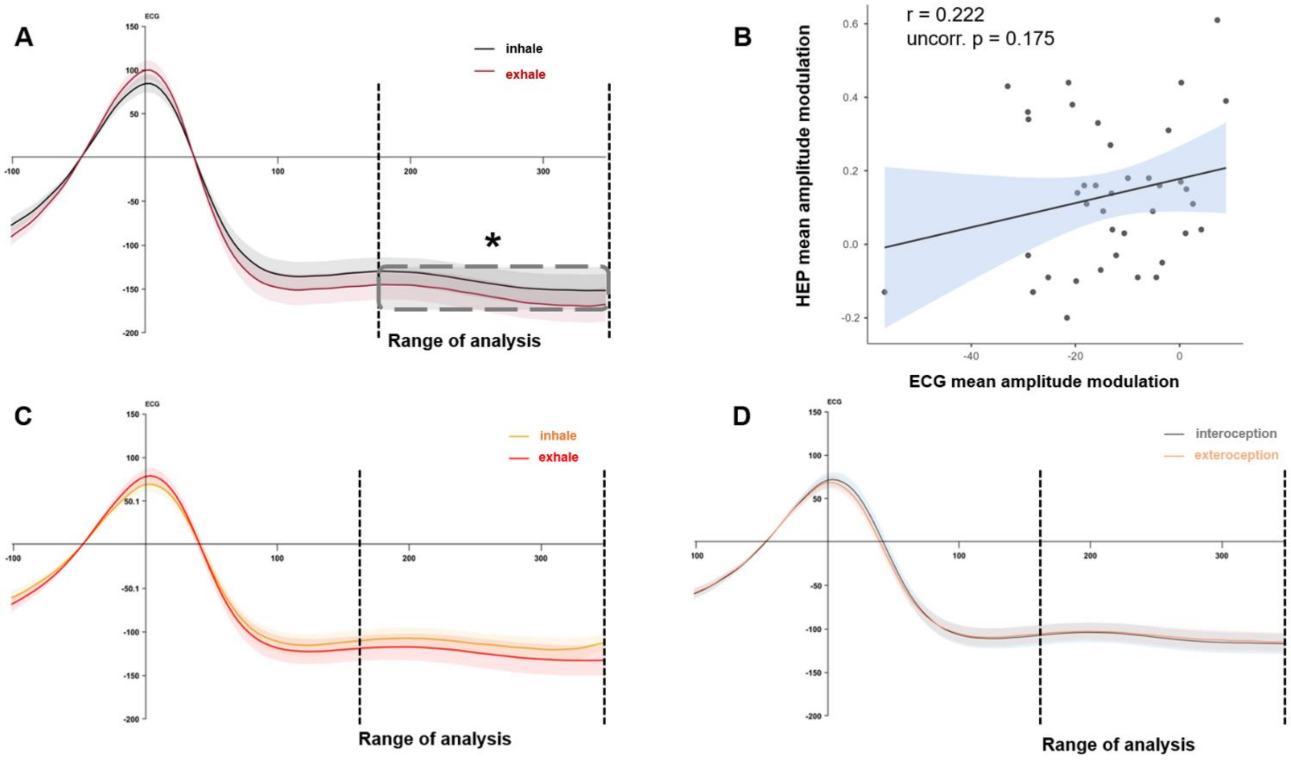


**Supplementary Figure 1.** (A) Grand average ECG signal, time-locked to the T-peak, for the inhale (black) and the exhale (dark red) phases at rest. The dotted lines represent the time window of interest used for statistical analyses (180-350 ms after the T-peak). The grey rectangle highlights the interval where significant differences were observed (180–347 ms after the T-peak). (B) Scatter plot of the linear relationship between the mean HEP amplitude changes and the mean ECG amplitude changes (ΔECG) among respiratory phases. Mean HEP amplitude changes were computed by aggregating the following channels: FC1, FC2, FC3, Cz, C1, C2, C3, C4, CPz, CP1, CP2, CP3, CP4, Pz, P1, P3, P4, POz. (C) Grand average ECG signal, time-locked to the T-peak, for the inhale (orange) and the exhale (light red) phases during the Interoceptive Condition of the HBD task. (D) Grand average ECG signal, time-locked to the T-peak, for the Interoceptive (grey) and Exteroceptive (yellow) conditions of the HBD task.
